# Antibody-mediated delivery of LIGHT to the tumor boosts Natural Killer cells and delays tumor progression

**DOI:** 10.1101/2020.05.08.084277

**Authors:** Marco Stringhini, Jacqueline Mock, Vanessa Fontana, Patrizia Murer, Dario Neri

## Abstract

**Background:** LIGHT is a member of the TNF superfamily, which has been claimed to mediate anti-tumor activity on the basis of cancer cures observed in immunocompetent mice bearing transgenic LIGHT-expressing tumors. The preclinical development of a LIGHT-based therapeutic has been hindered by the lack of functional stability exhibited by this protein.

**Methods:** Here, we describe the cloning, expression and characterization of five antibody-LIGHT fusion proteins, directed against the alternatively-spliced EDA domain of fibronectin, a conserved tumor-associated antigen.

**Results:** Among the five tested formats, only the sequential fusion of the F8 antibody in single-chain diabody format, followed by the LIGHT homotrimer expressed as a single polypeptide, yielded a protein (termed “F8-LIGHT”) which was not prone to aggregation. A quantitative biodistribution analysis in tumor bearing mice, using radioiodinated protein preparations, confirmed that F8-LIGHT was able to preferentially accumulate at the tumor site, with a tumor-to-blood ratio of ca. five to one twenty-four hours after intravenous administration. Tumor therapy experiments, performed in two murine tumor models (CT26 and WEHI-164), featuring different levels of lymphocyte infiltration into the neoplastic mass, revealed that F8-LiGHT could significantly reduce tumor cell growth and was more potent than a similar fusion protein (KSF-LIGHT), directed against hen egg lysozyme and serving as negative control of irrelevant specificity in the mouse. At a mechanistic level, the activity of F8-LiGHT was mainly due to an intratumoral expansion of Natural Killer cells, whereas there was no evidence of expansion of CD8+ T cells, neither in the tumor, nor in draining lymph nodes.

**Conclusion:** We developed a novel recombinant LIGHT fusion protein, able to accumulate at the site of disease and which displayed anti-tumor activity in two mouse models of cancer.

## Background

Immunotherapy of cancer is gaining momentum based on the realization that a subset of patients may enjoy durable responses upon treatment with immunotherapeutic agents [1]. Immune checkpoint inhibitors (e.g., antibodies directed against PD-1, PD-L1 and CTLA-4) have shown activity against different types of cancer and are widely used in the clinical practice [2–4]. Cytokines represent a complementary class of therapeutic proteins which modulate the activity of the immune system and engineered cytokine products are increasingly being used (alone or in combination) for oncological applications [5–7]. Cytokines are typically active in patients at very low doses (often at less than 1 milligram) and have a narrow therapeutic window. Research efforts have been dedicated to the development of more selective types of cytokine-based therapeutics with improved activity and/or safety.

Tumor-targeting antibody-cytokine fusions (also called “immunocytokines”) represent a promising class of anticancer agents and some of these products have moved to advanced clinical trials [8]. A preferential accumulation of the cytokine payload in the neoplastic mass may help boosting the activity of tumor-resident T cells and Natural Killer (NK) cells locally [9], resulting in lower systemic toxicity [10, 11]. In a number of comparative preclinical studies, tumor-homing immunocytokines were substantially more active than fusions based on antibodies of irrelevant specificity, even though this difference may be less drastic for long-lived IgG-based products [12–17].

LIGHT (which stands for homologous to **L**ymphotoxin, exhibits **I**nducible expression and competes with HSV **G**lycoprotein D for binding to **H**erpesvirus entry mediator, a receptor expressed on **T** lymphocytes) is a member of the tumor necrosis factor (TNF) superfamily expressed on a number of immune cells, including immature dendritic cells and activated T lymphocytes [18]. The membrane-anchored form of LIGHT can be cleaved by proteases between residues L81 and I82, resulting in a functional soluble form [19]. This cytokine binds to two receptors: HVEM (expressed on T cells, NK cells and dendritic cells) and LTβR (expressed mainly on non-lymphoid cells) [20]. Upon homotrimerization and interaction with HVEM, LIGHT activates the NF-κB pathway leading to activation and stimulation of target lymphocytes, whereas signaling through LTβR leads to expression of chemokines and adhesion molecules involved in lymphoid organs organization and maintenance [18, 21, 22]. Transgenic expression of LIGHT on tumor cells has been shown to mediate a potent anti-tumor effect in mice [19, 23]. In two independent studies, LIGHT expression resulted in a massive increase of CD8+ T cells infiltration, which played a central role in the tumor rejection process, as demonstrated by depletion experiments [19, 23].

A fusion of murine LIGHT with the F8 antibody (specific to the alternatively-spliced EDA domain of fibronectin, a conserved tumor-associated antigen) has previously been described, using the antibody in scFv format and relying on the ability of LIGHT to form stable non-covalent homotrimers [24]. The resulting fusion protein was homogeneous in biochemical characterization assays and maintained binding ability *in vitro*. However, in contrast to the results obtained with fusions of the F8 antibody with both murine and human TNF (which showed selective uptake at the tumor site with excellent biodistribution results), the fusion of scFv(F8) and murine LIGHT exhibited a poor uptake at the tumor site and a rapid clearance from the body [24]. It has previously been reported that members of the TNF superfamily may gain stability when expressed as a single polypeptide, with linkers connecting the three monomeric subunits [25, 26].

Here, we describe the cloning, expression and characterization of five novel fusion proteins, featuring murine LIGHT expressed as a single polypeptide (comprising the three monomeric subunits) as cytokine payload and the F8 antibody in different formats as tumor-targeting agent. Out of the five fusion proteins that were produced and purified, only the use of the F8 antibody in single-chain diabody format resulted in a product with adequate biochemical and immunological properties. The fusion protein preferentially accumulated in tumor lesions and mediated a potent anticancer activity, which mainly depended on Natural Killer cells and which was not potentiated by PD-1 blockade.

## Methods

### Cell lines and animal models

CT26 colon carcinoma (ATCC CRL-2638), WEHI-164 fibrosarcoma (ATCC CRL-1751), F9 teratocarcinoma (ATCC CRL-1720) and HT-29 (ATCC HTB-38) human adenocarcinoma cells were obtained from ATCC, CHO-S cells from Invitrogen. Cells were handled according to supplier’s protocol and maintained as cryopreserved aliquots in liquid nitrogen. Authentication including check of post-freeze viability, growth properties and morphology, test for mycoplasma contamination, isoenzyme assay and sterility test were performed by the cell bank before shipment. Tumor cell lines were kept in culture no longer than 3 weeks, CHO-S cells no longer than 4 weeks. Eight-weeks-old female BALB/c or 129/Sv mice were obtained from Janvier (France). All animal experiments were performed under a project license granted by the Veterinäramt des Kantons Zürich, Switzerland (04/2018).

### Cloning, expression and biochemical characterization of fusion proteins

The DNA sequence encoding murine LIGHT extracellular domain (amino acids 87-239) in a single-chain format, in which three LIGHT subunits were genetically linked together by a Glycine codon, including a C-terminal (SSSSG)_3_-linker, was purchased from Eurofins genomics. The LIGHT gene was fused by PCR assembly to the N-terminal sequence of various formats of the F8 antibody via its 15 amino acids linker. The resulting genes were cloned into the mammalian vectors pcDNA3.1+ (for F8 in scFv-Fc and diabody formats) or pMM137 (for F8 in IgG format) by restriction enzymes digestion and ligation, followed by amplification in TG1 electrocompetent *E. coli* bacteria. pMM137 was kindly provided by Philochem AG and has been described elsewhere [27]. Fusion proteins were produced in CHO-S by transient gene expression as already described [28, 29]. Both “low density” (LD) [28] and “high density” (HD) [29] protocols were used. Proteins were purified to homogeneity by protein A affinity chromatography and characterized by size exclusion chromatography on a Äkta Pure FPLC system (GE Healthcare) with a Superdex S200 10/300 increase column (GE Healthcare) and by SDS-PAGE.

### Mass spectrometry analysis of F8-LIGHT

The fusion protein was treated with glycerol free PNGase F (NEB, P0705S) in non-denaturing reaction conditions, as indicated by the manufacturer, to remove N-linked glycans. The resulting protein was analyzed by Liquid chromatography-Mass spectrometry on a Waters Xevo G2-X2 Qtof instrument coupled to a Waters Acquity UPLC H-class system, using a 2.1 x 50 mm Acquity BEH300 C4 1.7 μm column (Waters).

### Functional in vitro characterization of F8-LIGHT

Binding of the F8 moiety to its cognate antigen was tested by ELISA. Briefly, wells of a F96 Maxisorp nunc-immuno plate (Thermofisher) were coated with EDA and blocked with 2% milk powder in PBS. F8-LIGHT and positive control F8 in small immune protein format (SIPF8) were added at different concentrations in wells, followed by incubation with polyclonal antibodies against human kappa light chains (Dako, A0192) and by HRP-conjugated protein A (GE Healthcare, NA9120V). Colorimetric reaction was started by adding BM Blue POD substrate (Roche, 11442066001) and quenched with 1M H_2_SO_4_. Absorbance at 450 nm was measured with a Spectra Max Paradigm multimode plate reader (Molecular Devices). *In vitro* activity of the LIGHT moiety was tested with a cytotoxicity assay on HT-29 cells, which have been shown to be sensitive to LIGHT in the presence of human interferon gamma (hIFNγ) [31]. Briefly, F8-LIGHT and recombinant hIFNγ (Biolegend, 570202) at given concentrations were added to wells containing 40000 HT-29 cells in McCoy’s 5A (Modified) Medium (Thermofisher, 16600082) supplemented with 10% fetal bovine serum (Thermofisher, 10270106) and 1x Antibiotic-Antimycotic (Thermofisher, 15240062). After incubation for 72 hours at 37°C, 5% CO_2_, cells were detached using Trypsin-EDTA solution (Thermofisher, 25200056), re-suspended in PBS containing 0.5% BSA and 2mM EDTA and stained with 7-AAD (Biolegend, 420404) for 5’ at 4°C. Percentage of living cells was determined by Flow cytometry analysis using a CytoFLEX S (Beckman Coulter). Data were analyzed with FlowJo software (FlowJo, LLC, version 10).

### Quantitative biodistribution

*In vivo* targeting ability of F8-LIGHT was determined by quantitative biodistribution as already described [32]. Briefly, 1×10^7^ F9 cells were subcutaneously injected in the right flank of 129/Sv mice. Tumor size was determined daily using the formula: ½ x (major diameter) x (minor diameter)^2^. When tumors reached a volume of 100-300 mm^3^, fusion proteins were labeled with Iodine-125, using Chloramine T. Radiolabeled protein was purified by size exclusion chromatography (PD-10 column, GE Healthcare, 17-0851-01) and injected in the lateral tail vein of tumor-bearing mice. Animals were sacrificed after 24 hours. Organs and tumors were harvested, weighted and radioactivity was determined using a Packard Cobra II Gamma Counter (GMI).

### Tumor therapy experiments

Anti-tumor activity of F8-LIGHT was tested in two syngeneic murine models of cancer. CT26 and WEHI-164 tumors were implanted in the right flank of BALB/c mice by subcutaneous injection of 3×10^6^ resp. 2.5×10^6^ cells. Treatments were started when tumors reached a volume of about 100 mm^3^. The indicated dose of immunocytokines or the corresponding volume of saline, were administered by intravenous injection in the lateral tail vein, every other day for a total of three injections. The immune-checkpoint inhibitor anti-PD-1 (BioXCell, clone 29F.1A12) was administered in the same way, at alternate days. Mice were inspected daily. Body weight was monitored, tumor was measured with a caliper and tumor size was determined using the formula: ½ x (major diameter) x (minor diameter)^2^. Mice with ulcerated hemorrhagic tumor, or with tumor bigger than 1500 mm^3^ where euthanized.

### Antibody and reagents for Flow Cytometry

Fluorophore-conjugated antibodies against CD3 (clone 17A2), CD4 (clone GK1.5), CD8 (clone 53-6.7), NK1.1 (clone PK136), I-A/I-E (clone M5/114.15.2), CD62L (clone MEL-14), CD44 (clone IM7), and FoxP3 (clone MF-14), as well as 7-AAD and Zombie Red viability dies were all purchased from BioLegend. AH1-loaded, PE-conjugated H-2L^d^ tetramers were obtained as already described [33].

### Analysis of immune infiltrates

Immune infiltrates were analyzed in tumors and tumor-draining lymph nodes of mice bearing WEHI-164 sarcoma, 48 hours after the last injection of either saline or F8-LIGHT. Mice were euthanized and tumors, right axillary and right inguinal lymph nodes were harvested. Tumors were cut into small fragments and incubated in an orbital shaker at 37°C for 30’ in RPMI 1640 Medium (Thermofisher, 21875034) containing 1x Antibiotic-Antimycotic (Thermofisher, 15240062), 1 mg/mL Collagenase II (Thermofisher, 17101015) and 0.1 mg/mL DNAse I (Roche, 10104159001). Tumor cells suspensions were passed through a 70μm cell strainer (Corning) and treated with Red Blood Cells Lysis buffer (Biolegend, 420301) following supplier’s recommendations. Lymph nodes were harvested and smashed on a 70μm cell strainer (Corning) using the back of a syringe plunger. The resulting tumor and lymph nodes single cell suspensions were washed in PBS, before incubation with staining reagents. Where appropriate, cells were stained with Zombie Red dye (diluted 1:500 in PBS) for 15’ at room temperature, followed by staining with antibodies and tetramers in FACS buffer (0.5% BSA, 2mM EDTA in PBS) for 30’ at 4°C. Cells, which were not stained with Zombie Red, were stained with 7-AAD (diluted 1:100 in FACS buffer) for 5’ at 4°C. Intracellular staining with antibodies against FoxP3 was performed using eBioscience^™^ Foxp3 / Transcription

Factor Staining Buffer Set (Thermofisher), following supplier’s protocol. Samples were analysed with CytoFLEX S (Beckman Coulter) and data were processed using FlowJo software (FlowJo, LLC, version 10). A detailed description of the gating strategies can be found in Additional files (Supp. Fig. 5-7). Total of living cells in the tumor was calculated by subtracting dead cells and debris from the total number of recorded events.

### Data analysis

Data were analyzed using Prism 7.0 (GraphPad Software, Inc.). Statistical significances of tumor therapy and flow cytometry data were determined with a regular two-way ANOVA test with Bonferroni post-test correction and with an unpaired, two-tailed t-test, respectively. Data represent means ± SEM. P < 0.05 was considered statistically significant.

## Results

### Expression and characterization of F8-LIGHT fusion proteins

LIGHT is a homotrimeric protein, that can activate HVEM and LTβ receptors [13, 17] (Figure 1A). We cloned, expressed and characterized five different versions of the F8 antibody, fused to murine homotrimeric LIGHT expressed as a single polypeptide. Specifically, we fused LIGHT at the C-terminus of the heavy or of the light chain of the F8 antibody in human IgG1 format, at the C-terminus of the F8 antibody in human scFv-Fc format and at the C-terminus of the F8 antibody in human single-chain diabody and diabody format. (Figure 1B). All products could be purified to homogeneity by protein A chromatography, as shown by SDS-PAGE analysis. However, a comparative evaluation of size-exclusion chromatography profiles revealed that only the single-chain diabody-based product (termed F8-LIGHT), ran as a single peak in gel filtration (Figure 1B), showed a single band in SDS-PAGE and a single peak in mass spectrometry, after PNGase F treatment (Figure 1C). F8-LIGHT was able to bind its cognate target antigen with high affinity *in vitro*, as shown by ELISA (Figure 1E) and was able to induce death of HT-29 cells in the presence of interferon gamma (Figure 1F). When the cytotoxicity experiment on HT-29 cells was repeated with commercially available recombinant murine LIGHT as control, F8-LIGHT showed a higher activity, possibly as a result of the increased stability of the LIGHT moiety expressed as a covalently-linked homotrimer [25, 26] (Additional files, Supp. Fig. 1). Dissolved in saline, the product was stable for months both at 4°C and at −80°C, even after repeated cycles of freeze-and-thaw (Additional files, Supp. Fig. 2).

**Figure 1.**
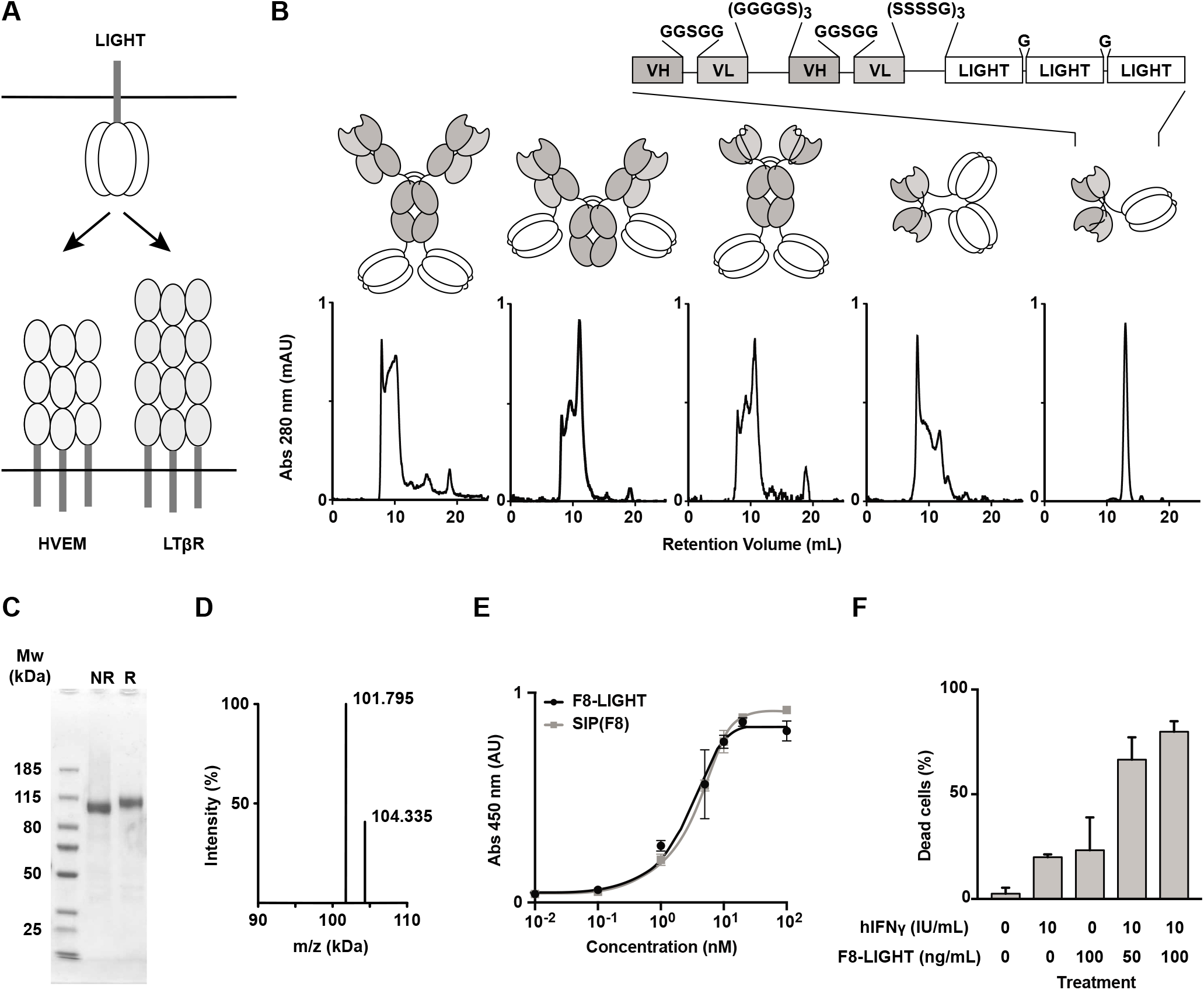
*In vitro* characterization of fusion proteins. **A**, schematic representation of membrane-anchored LIGHT and cognate receptors HVEM and LTβR. **B**, schematic representation of five LIGHT-based fusion proteins, with respective size exclusion chromatography profiles. The homotrimeric form of LIGHT expressed as a single polypeptide chain was fused to the C-terminus of (from left to right): the heavy chain of F8 in IgG format, the light chain of F8 in IgG format, the F8 in scFv-Fc format, the F8 in diabody format and the the F8 in single-chain diabody format (F8-LIGHT). Detailed linear structure of F8-LIGHT is highlighted. **C-D**, biochemical characterization of F8-LIGHT including SDS-PAGE of F8-LIGHT under non-reducing (NR) and reducing (R) conditions (**C**) and mass spectrometry profile of PNGase F-treated F8-LIGHT (calculated mass = 101791 Da) (**D**). **E**, binding of titrated concentrations of F8-LIGHT and positive control SIP(F8) to immobilized target antigen EDA, measured by ELISA. **F**, activity of F8-LIGHT, measured by a cytotoxicity assay on HT-29 cells in the presence of human Interferon gamma (hIFNγ). Reported concentrations are based on the molecular weight of the LIGHT part of the molecule alone. 7-AAD positive dead cells were detected by Flow Cytometry.

### F8-LIGHT selectively accumulate at tumor site *in vivo*

In order to test the *in vivo* targeting ability of our product, we performed a quantitative biodistribution study by intravenous injection of ^125^I-labeled F8-LIGHT. We used LIGHT linked to the KSF antibody (specific to hen egg lysozyme) as negative control of identical format (Figure 2). After 24h, about 4.5% of the injected dose of F8-LIGHT per gram of tissue was found in the tumor, with a tumor-to-blood ratio of 4.9. Similar to what previously reported for other antibody-cytokine fusion proteins, the transfection protocol used for transient gene expression procedures had an impact on biodistribution results [34, 35] (Additional files, Supp. Fig. 3).

**Figure 2.**
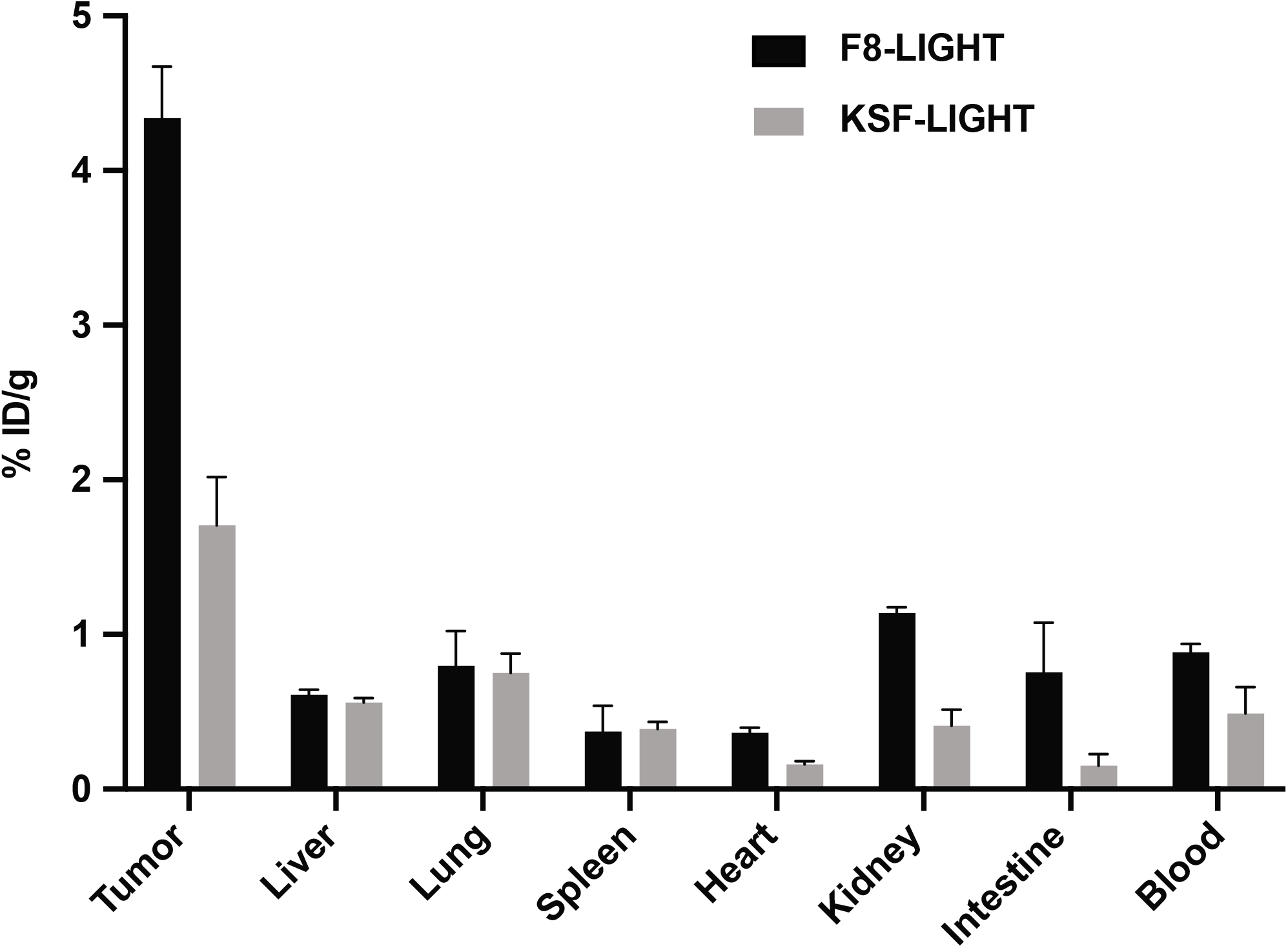
Tumor targeting of F8-LIGHT *in vivo*. Biodistribution experiment in 129/Sv mice bearing F9 tumor. Radioiodinated F8-LIGHT and KSF-LIGHT (used as negative control) were injected in the lateral tail vein. Accumulation of the fusion proteins in tumor and healthy organs after 24 hours was calculated as percentage of injected dose per gram of tissue (% ID/g). Column represent means ± SEM, n = 4 per experimental group.

### F8-LIGHT delay progression of established murine tumors

To evaluate the anti-tumor activity of F8-LIGHT, we performed a first therapy experiment in BALB/c mice bearing subcutaneous CT26 murine colon carcinoma, since forced expression of LIGHT in this model had previously shown the ability to induce complete tumor regression [23]. Treatment was initiated when tumors had reached a volume of about 100 mm^3^ and consisted in intravenous injection of 100 μg F8-LIGHT every other day, for a total of three injections. Treatment with F8-LIGHT induced tumor growth retardation, whereas the KSF-LIGHT fusion protein used as negative control gave profiles similar to the ones obtained in the saline treatment group. As no toxicity had been observed (Figure 3A), the dose was increased to 300 μg/injection, which was still well tolerated but did not further improve anti-cancer activity (Figure 3B). CT26 is an immunologically “hot” murine tumor that exhibits a rich infiltrate of lymphocytes [30, 36]. Flow cytometry experiments on saline treated tumors showed that the proportion of T lymphocytes in CT26 tumors was higher than in other commonly studied BALB/c tumors, such as WEHI-164 sarcoma or C51 colon carcinoma (Additional files, Supp. Fig. 4A). Interestingly, also the percentage of CD8+ T cells specific to the AH1 rejection antigen was significantly higher in CT26, compared to the other models (Additional files, Supp. Fig. 4B). In order to study anticancer activity in a second immunocompetent mouse model, we treated WEHI-164 sarcomas and we included a combination treatment with PD-1 blockade. Also in this case, F8-LIGHT monotherapy significantly inhibited tumor-growth compared to saline. Combination treatment with an anti-PD-1 antibody led to tumor regression in all mice. Two out of five animals enjoyed a complete and durable remission, but a similar activity was also observed in the PD-1 blockade monotherapy group (Figure 3C).

**Figure 3.**
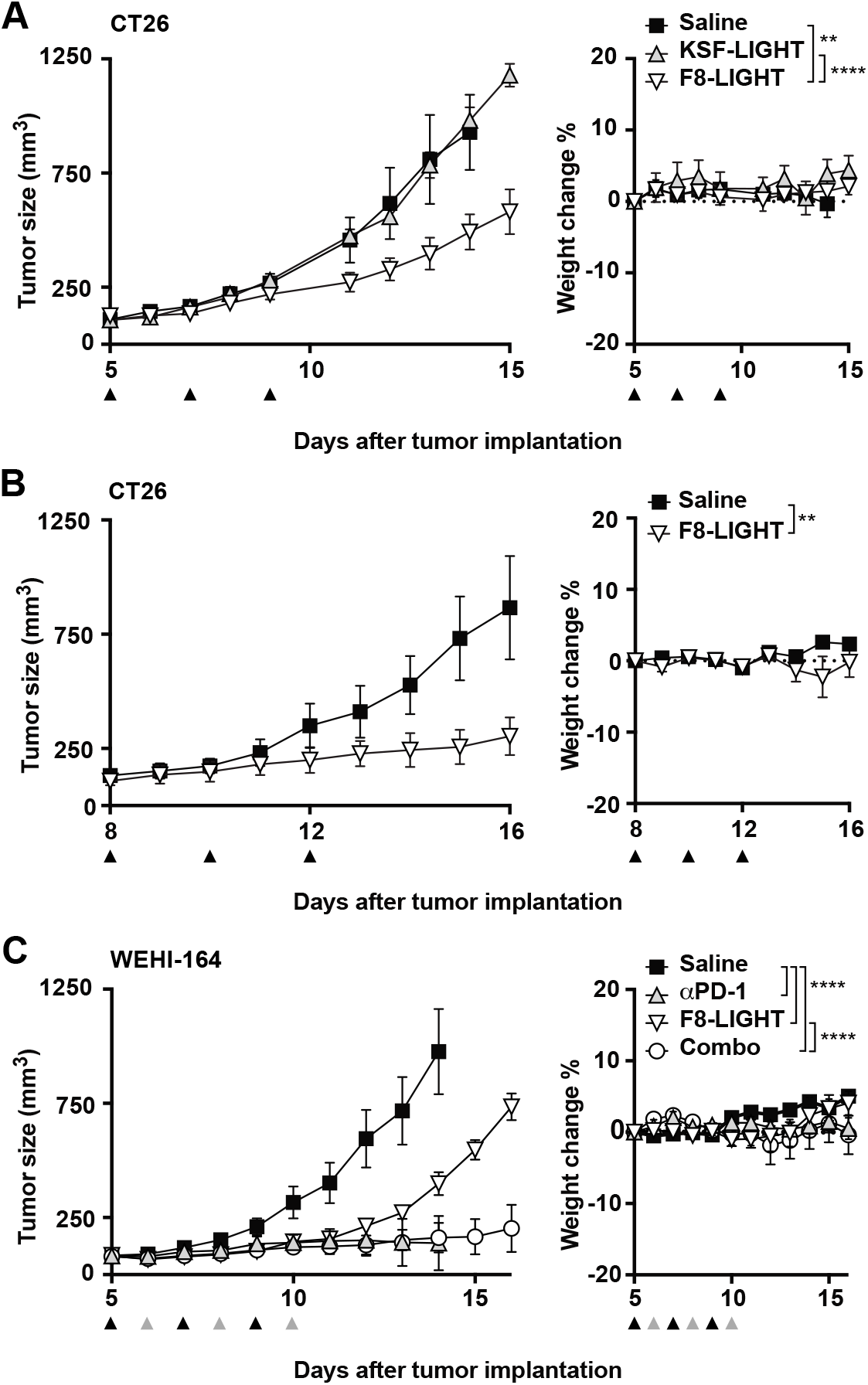
Therapy experiments. **A**, Therapy experiment in BALB/c mice bearing established CT26 tumor. 100 μg of F8-LIGHT or KSF-LIGHT were administered intravenously every other day, as indicated by the black arrows. **B**, as in **A**, but mice received 300 μg of F8-LIGHT. **C**, Therapy experiment in BALB/c mice bearing established WEHI-164 tumor. 300 μg of F8-LIGHT (black arrows), 200 μg of αPD-1 (grey arrows) or a combination of the two were administered as depicted in the figure. Data represent means ± SEM, n = 5 mice per experimental group. * = p < 0.05, ** = p < 0.01, *** p = < 0.001, **** = p < 0.0001 (regular two-way ANOVA test with Bonferroni post-test correction).

### F8-LIGHT treatment increase Natural Killer cells but not CD8+ T cells infiltration in the tumor

Since previous studies had reported the ability of LIGHT to attract CD8+ T cells into the neoplastic mass, we treated WEHI-164 bearing mice with F8-LIGHT or saline and analyzed tumors and tumor-draining lymph nodes (harvested 48 hours after the last injection) by flow cytometry. In contrast to reports obtained by the transgenic expression of LIGHT by tumor cells, we did not observe any increase of CD8+ T cells in tumors from the F8-LIGHT treatment group compared to saline (Figure 4A). Instead, we found a significant increase of tumor-infiltrating cells positive for the Natural Killer cells marker NK1.1 (Figure 4B). Within the NK1.1 positive population, we could distinguish three different subpopulations: a CD3-negative (conventional NK cells), a CD3-positive (NKT cells) and a CD3-intermediate, MHC class II-positive population, which was the most abundant one (Figure 4A). Inspection of the phenotype composition of the CD8+ T cells revealed an increase in the proportion of CD62L+CD44low naïve CD8+ T cells infiltrating LIGHT-treated tumors (Figure 4C). The same feature was observed for the AH1-specific CD8+ T cell population in the tumor, whereas in the draining lymph node the situation was reversed, with an increased proportion of CD62L-CD44+ effector AH1-specific T cells after F8-LIGHT treatment (Figure 4C). No significant difference among the different therapy groups could be found in terms of abundance of CD3+, CD4+, MHC class II+ and AH1-specific CD8+ cells. No difference in CD8-to-CD4 ratios was observed in tumors or lymph nodes (Figure 4A, D-E).

**Figure 4.**
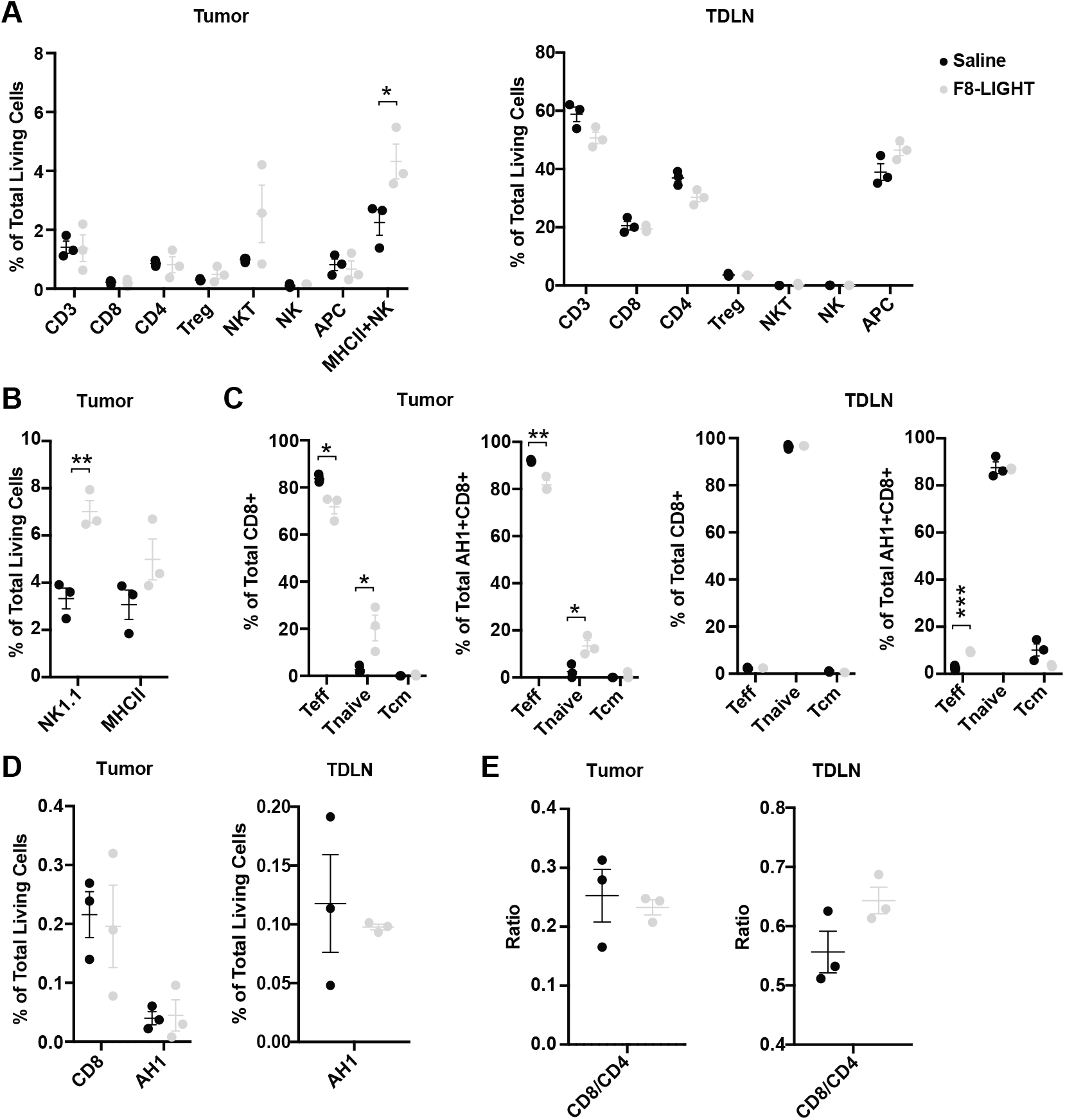
Analysis of immune infiltrate. Analysis of tumors and tumor-draining lymph nodes (TDLN) of BALB/c mice bearing WEHI-164 tumors, 48 hours after the third administration of F8-LIGHT or saline. **A**, lymphocytes infiltration in tumors and composition of TDLN. CD3 = CD3^+^ lymphocytes, CD8 = CD8+ T cells, CD4 = CD4+ T cells, T_reg_ = CD4+ regulatory T cells, NKT = Natural Killer T cells, NK = Natural Killer cells, APC = Antigen Presenting cells and MHCII+NK = MHC class II+ NK cells. **B**, total NK1.1-positive and MHC class II-positive lymphocytes infiltrating tumors. **C**, phenotype of CD8+ T cells and of AH1-specific CD8^+^ T cells in tumors and TDLN, based on expression of the markers CD44 and CD62L. T_eff_ = effector T cells, T_naive_ = naïve T cells, T_cm_ = central memory T cells. **D**, AH1-specific CD8+ T cells in tumors and TDLN. **E**, CD8+ T cells:CD4+ T cells ratio in tumor and TDLN. Data represent individual mice, with means ± SEM, n = 3 mice per experimental group. * = p < 0.05, ** = p < 0.01, *** p = < 0.001 (unpaired t-test).

F8-LIGHT treatment mediated a tumor growth retardation, but the proportion of total living cells within the neoplastic mass (determined by 7-AAD staining on total events at time of analysis) was significantly higher in the F8-LIGHT group compared to saline, while leukocyte levels were similar (Figure 5A-B). When trying to identify the nature of living cells within the F8-LIGHT-treated neoplastic mass, we observed fewer tumor cells compared to the saline group, accompanied by the emergence of an abundant population of FSC-low and SSC-low particles (which we called SLE, for “small living events”) (Figure 5C). These SLEs stained negative for 7-AAD, CD3, CD4, CD8, NK1.1 and MHC class II (Additional files, Supp. Fig. 6).

**Figure 5.**
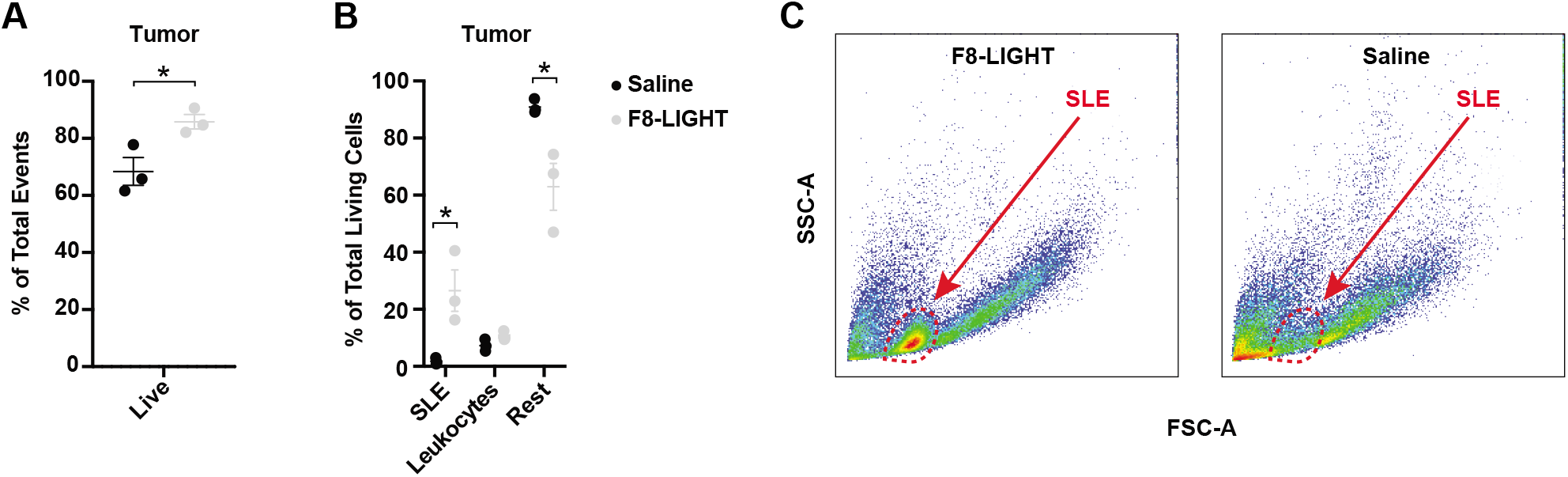
Analysis of tumor composition. WEHI-164 tumors analysis 48 hours after the third administration of F8-LIGHT or saline **A**, fraction of living cells among total events recorded. **B**, composition of living cells in the tumor. “Leukocytes” represent the sum of all T cells, NK cells, Antigen Presenting cells and Granulocytes, “Rest” represent the remaining living cells, after subtracting SLE and Leukocytes from the total number of living cells. SLE = “Small Living Events” **C**, representative analysis of tumor cells suspensions from single mice treated with F8-LIGHT or saline. SSC-A = side scatter area, FSC-A = forward scatter area. Data represent individual mice, with means ± SEM, n = 3 mice per experimental group. * = p < 0.05 (unpaired t-test).

## Discussion

We have generated a novel fusion protein (termed F8-LIGHT), which was able to selectively deliver murine LIGHT to solid tumors, thanks to the binding properties of the F8 antibody, specific to the EDA domain of fibronectin. This immunocytokine product was well-behaved in biochemical assays and fully active *in vitro*, unlike other formats featuring murine LIGHT as module, described in this article or in previous studies [37, 38]. F8-LIGHT induced a specific tumor-growth retardation, that was not observed for a similar fusion protein used as negative control of irrelevant specificity in the mouse (KSF-LIGHT). In spite of its potent anti-cancer activity, F8-LIGHT could not cure CT26 or WEHI-164 tumors in mice, when used as a single agent.

The strongest evidence linking LIGHT to an anti-cancer activity had emerged from the use of tumor cells, which had been engineered to express large quantities of LIGHT and which were completely rejected by immunocompetent mice in a process that relied on CD8+ T lymphocytes [19, 23, 39]. In those studies, it is difficult to control and measure the amounts of LIGHT expressed by the tumor cells. As a consequence, even the use of tumor-homing LIGHT fusion proteins may not reach comparable *in vivo* concentrations of cytokine at the site of disease. A further difference between the two experimental set ups may relate to the fact that F8-LIGHT anchored the cytokine payload on the tumor extracellular matrix, which contained the EDA domain of fibronectin, while transgenic tumor cells may display LIGHT on their surface [19, 23, 32]. The antibody-based delivery of cytokines to components of the modified extracellular matrix (e.g., splice variants of fibronectin or of tenascin-C) has been shown to be potently active for other immunomodulatory agents, including interleukin-2, interleukin-12 and tumor necrosis factor [11, 14, 40].

Although many studies have focused on the effect of LIGHT on CD8+ lymphocytes [19, 41], Fan Z. et al. demonstrated the essential contribution of NK cells at an early phase of the LIGHT-induced anti-tumor response [39]. They showed that LIGHT was capable of activating and inducing proliferation of NK cells and that the interferon gamma produced by these NK cells contributed to the subsequent activation of CD8+ cells. Treatment of tumor-bearing mice with F8-LIGHT led to an expansion of intratumoral NK cells, but not of CD8+ T cells. The inability to potently activate the T cell compartment within the neoplastic mass was confirmed by a lack of synergy with murine PD-1 blockade (Figure 3).

LIGHT is a member of the TNF superfamily. Our group and other researchers have previously studied the possibility to use tumor-homing antibodies for the selective delivery of TNF and related homotrimeric proteins [17, 24, 25, 40, 42]. TNF fusions exhibit an extremely selective tumor-targeting performance, possibly due to the vasoactive properties of their cytokine payload, while other superfamily members are difficult to deliver [24]. Besides falling in the second category in term of tumor targeting ability, murine LIGHT has also been reported to be difficult to express as a recombinant [37, 38, 43]. In our hands, LIGHT showed a selective accumulation into solid tumors only after extensive engineering of the protein format and of gene expression methods (Figure 1 and Additional files, Supp. Fig. 3). Similar features have recently been reported for other cytokine payloads, including interleukin-12 and interleukin-15 [34, 44, 45].

An EGFR-targeted version of a mutant human LIGHT (EGFR-LIGHT) has recently been described [38]. The fusion protein cross-reacted with murine HVEM and LTβ receptors, as shown by FACS analysis, and exhibited a potent antitumor activity in mice bearing small malignant lesions, but had limited effect in mice bearing more advanced tumors. Unlike what we observed for F8-LIGHT, EGFR-LIGHT treatment induced a strong infiltration of CD8+ T cells in the tumor mass. In a tumor model with a low natural level of immune infiltration, which did not respond to treatment with anti-PD-L1 or EGFR-LIGHT, combination with the two agents led to complete tumor eradication [38].

Experimental tumors and neoplastic masses in patients can be defined as “hot” and as “cold”, based on the relative density of lymphocyte infiltration. The CT26 model, used in this study and in many other investigations, is considered a “hot” tumor, which readily responds to immunotherapy [30, 36]. By contrast, WEHI-164 fibrosarcoma (also used in our study) exhibits a lower level of lymphocyte infiltration (Additional files, Supp. Fig. 4). Both models grow in BALB/c mice and in both models the antigenic peptide AH1 (derived from the gp70 envelope protein of the murine leukemia virus) can play a role for the tumor-rejection process [46]. Aberrantly-expressed antigens (including retroviral gene products) can act as dominant tumor-rejection antigens in some settings [47]. Unlike what we had previously observed for other antibody-cytokine products [11, 33, 40], treatment with F8-LIGHT did not substantially alter the tumor density of AH1-specific CD8+ T cells (Figure 4).

## Conclusion

A fully human analogue of F8-LIGHT may represent a useful tool for the *in vivo* boosting of NK cell activity, but this cytokine payload may be suboptimal for a selective activation of tumor-specific CD8+ T cells. The EDA domain of fibronectin represents an ideal target for preclinical studies and for clinical translational activities, as the antigen is highly conserved from mouse to man. EDA is expressed in the majority of solid and hematological malignancies [48, 49], while being virtually undetectable in normal adult tissues [50].

## Declarations

## Acknowledgements

D. N. gratefully acknowledges ETH Zürich, the Swiss National Science Foundation (Grant Nr. 310030_182003/1) and the European Research Council (ERC, under the European Union’s Horizon 2020 research and innovation program, grant agreement 670603) for financial support.

## Authors’ contributions

D.N. and M.S. planned and designed the study. M.S. performed the experiments. J.M. and P.M. helped with the biodistribution experiments. V.F. performed some experiments under the supervision of M.S. M.S. prepared the figures. D.N. and M.S. wrote the manuscript.

## Availability of data and material

Data and material are presented in the main text and supplementary information, raw data are available from the corresponding author on reasonable request.

## Ethics approval and consent to participate

Mice experiments were conducted under the ethical approval of the Veterinäramt des Kantons Zürich, Switzerland (License 04/2018).

## Abbreviations

PD-1: Programmed cell death protein 1
PD-L1: Programmed death-ligand 1
CTLA-4: Cytotoxic T-lymphocytes-associated protein 4
NK: Natural killer cells
LIGHT: Lymphotoxin, exhibits inducible expression and competes with HSV glycoprotein D for binding to herpesvirus entry mediator, a receptor expressed on T lymphocytes
TNF: Tumor necrosis factor
HVEM: Herpesvirus entry mediator
LTβR: Lymphotoxin beta receptor
NF-κB: Nuclear factor “kappa-light-chain-enhancer” of activated B cells
IFNγ: Interferon gamma
EGFR: Epidermal growth factor receptor.

