## Supplementary Information for "Antibody-mediated delivery of LIGHT to the tumor boosts Natural Killer cells and delays tumor progression"

### Additional file

#### F8-LIGHT amino acids sequence (Mw: 101790.95)

EVQLLESGGGLVQPGGSLRLSCAASGFTFSLFTMSWVRQAPGKGLEWVSAISGSGGSTYY
ADSVKGRFTISRDNSKNTLYLQMNSLRAEDTAVYYCAKSTHLYLFDYWGQGTLVTVSSGG
SGGEIVLTQSPGTLSLSPGERATLSCRASQSVSMPFLAWYQQKPGQAPRLLIYGASSRATG
IPDRFSGSGSGTDFTLTISRLEPEDFAVYYCQQMRGRPPTFGQGTKVEIKGGGGSGGGGS
GGGGSEVQLLESGGGLVQPGGSLRLSCAASGFTFSLFTMSWVRQAPGKGLEWVSAISGS
GGSTYYADSVKGRFTISRDNSKNTLYLQMNSLRAEDTAVYYCAKSTHLYLFDYWGQGTLV
TVSSGGSGGEIVLTQSPGTLSLSPGERATLSCRASQSVSMPFLAWYQQKPGQAPRLLIYGA
SSRATGIPDRFSGSGSGTDFTLTISRLEPEDFAVYYCQQMRGRPPTFGQGTKVEIKSSSSG
SSSSGSSSSGSHQANPAAHLTGANASLIGIGGPLLWETRLGLAFLRGLTYHDGALVTMEPG
YYYVYSKVQLSGVGCPQGLANGLPITHGLYKRTSRYPKELELLVSRRSPCGRANSSRVWW
DSSFLGGVVHLEAGEEVVVRVPGNRLVRPRDGTRSYFGAFMVGSHQANPAAHLTGANAS
LIGIGGPLLWETRLGLAFLRGLTYHDGALVTMEPGYYYVYSKVQLSGVGCPQGLANGLPITH
GLYKRTSRYPKELELLVSRRSPCGRANSSRVWWDSSFLGGVVHLEAGEEVVVRVPGNRL
VRPRDGTRSYFGAFMVGSHQANPAAHLTGANASLIGIGGPLLWETRLGLAFLRGLTYHDGA
LVTMEPGYYYVYSKVQLSGVGCPQGLANGLPITHGLYKRTSRYPKELELLVSRRSPCGRAN
SSRVWWDSSFLGGVVHLEAGEEVVVRVPGNRLVRPRDGTRSYFGAFMV

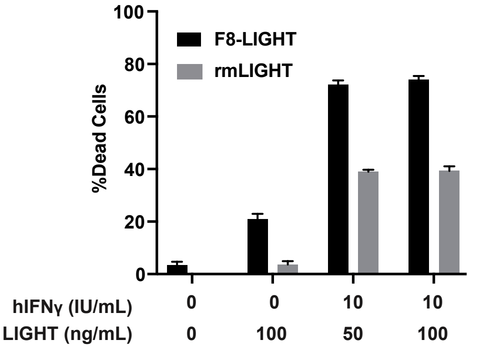


**Supplementary Figure 1.** ***In vitro*** **activity of F8-LIGHT versus recombinant murine LIGHT.**
Activity of F8-LIGHT and recombinant murine LIGHT (rmLIGHT, BioLegend) was compared in a cytotoxicity assay on HT-29 cells. hIFNγ = human Interferon gamma. Concentration are based on the molecular weight of the LIGHT monomer.

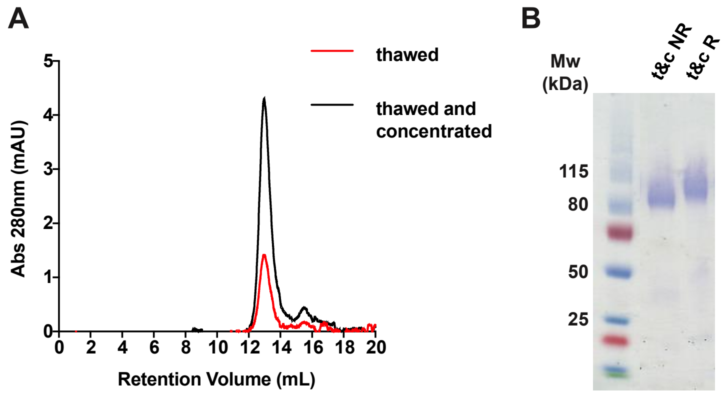


**Supplementary Figure 2. Stability of F8-LIGHT.**
Biochemical characterization of F8-LIGHT, after freeze and thaw and concentration by ultracentrifugation. **A**, size exclusion chromatography profile. **B**, SDS-PAGE under non-reducing (NR) and reducing (R) conditions, t&c = thawed and concentrated.

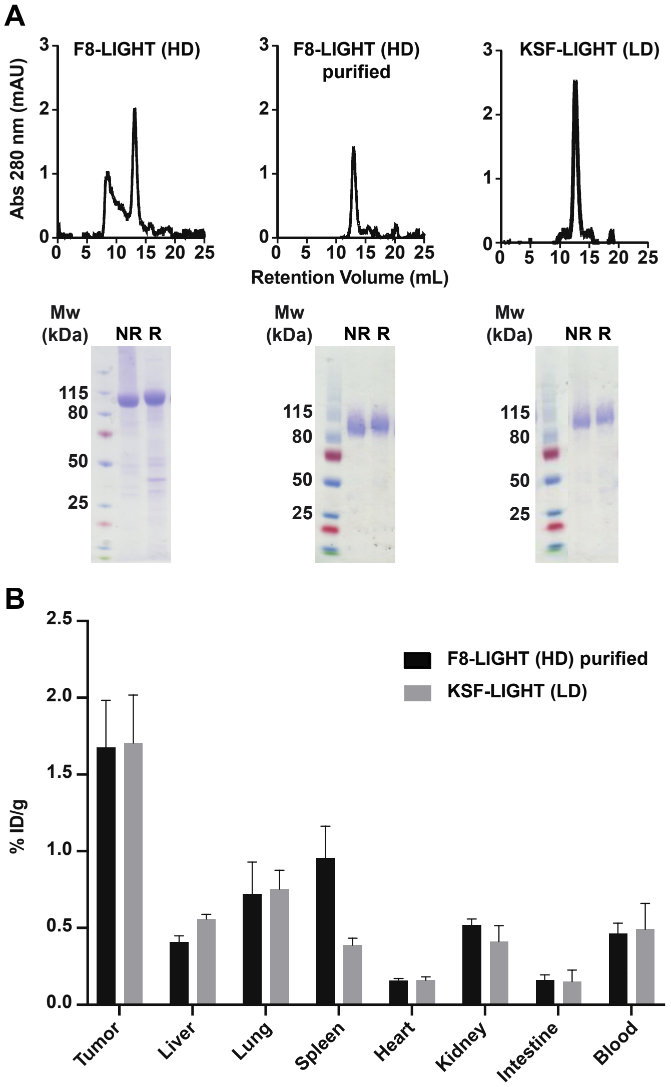


**Supplementary Figure 3. In vivo targeting of F8-LIGHT produced with the "High density" protocol.**
**A**, Biochemical characterization of F8-LIGHT produced with the HD protocol and KSF-LIGHT produced with the LD protocol, including size exclusion chromatography profiles and

SDS-PAGE under non-reducing (NR) and reducing (R) conditions of F8-LIGHT before and after purification by gel filtration (KSF-LIGHT, as F8-LIGHT, eluted as a single peak and did not need to be purified, when produced with the LD protocol). **B**, accumulation of radiolabelled preparations of purified F8-LIGHT (HD) and KSF-LIGHT (LD) in tumors and healthy organs of 129/Sv mice bearing F9 tumors, 24 hours after intravenous administration.

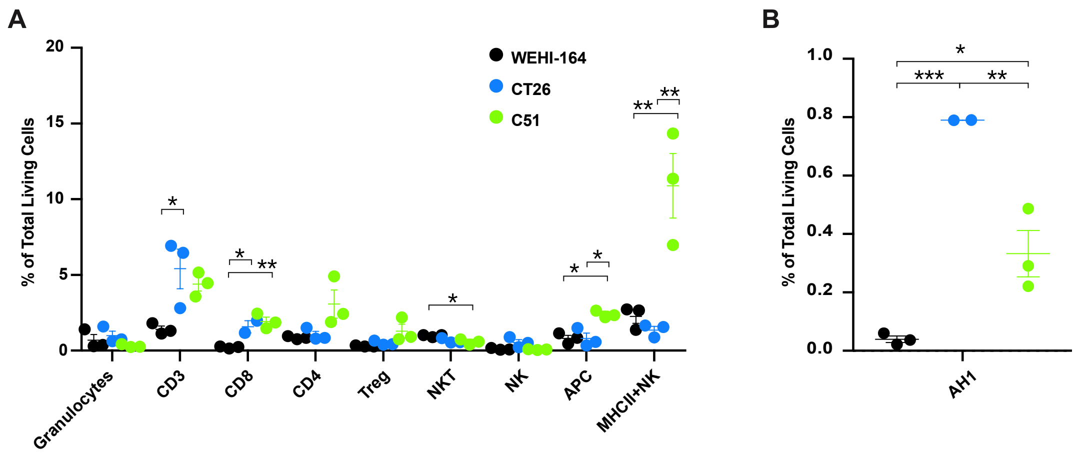


**Supplementary Figure 4. Analysis of leukocyte infiltrate in different BALB/c syngeneic tumor models.**

Flow Cytometry analysis of the immune infiltrate in BALB/c bearing WEHI-164, CT26 or C51 tumors, 48 hours after the third *i.v.* injection of saline (saline administered every other day starting when the tumor was about 100mm3). **A**, analysis of different population of leukocytes. **B**, analysis of tumor-infiltrating AH1-specific CD8+ T cells. Data represent means ± SEM, n = 3 mice per experimental group. * = $p < 0.05$, ** = $p < 0.01$, *** p = < 0.001 (regular one-way ANOVA test with Bonferroni post-test correction).

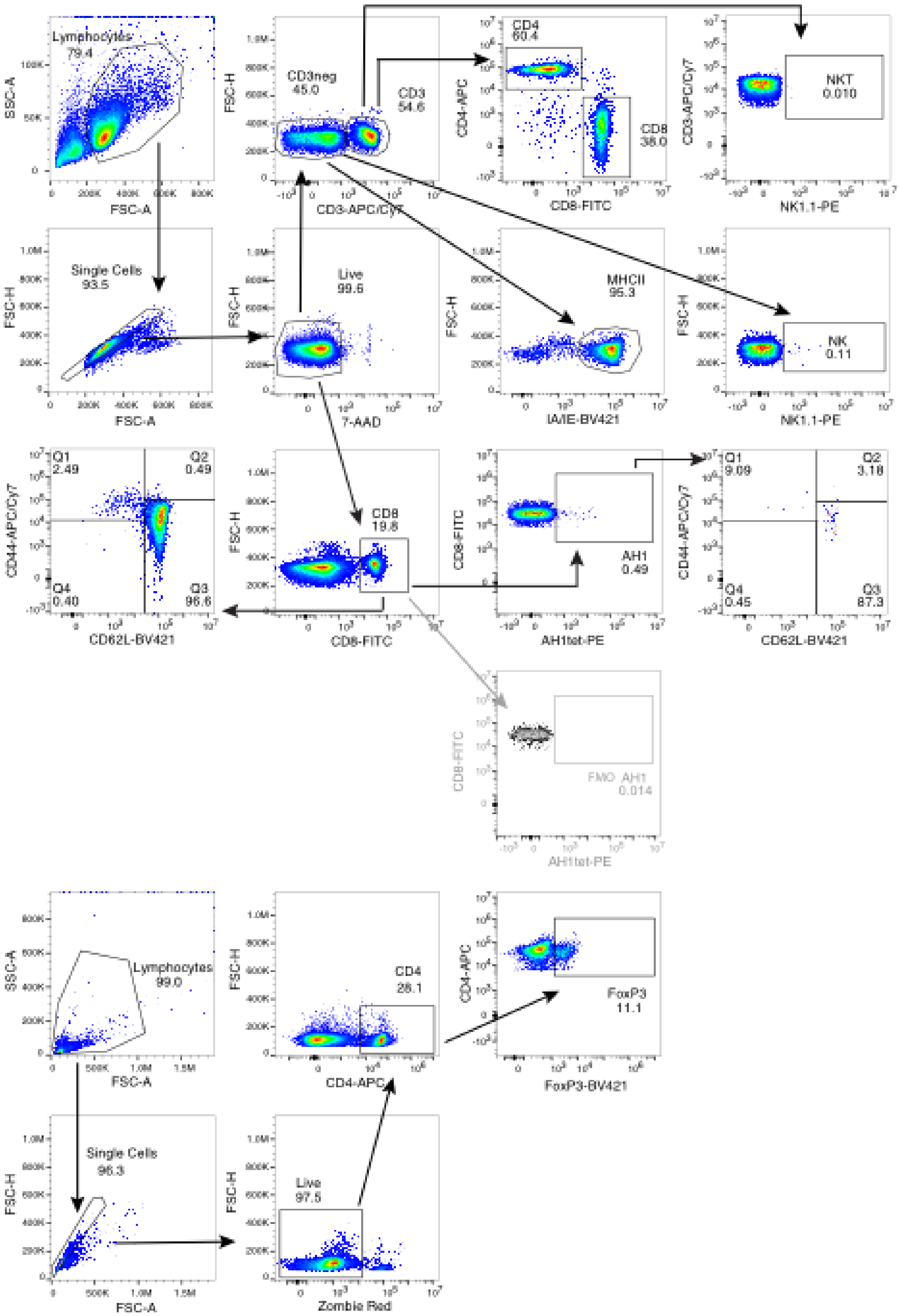


**Supplementary Figure 5. Gating strategy for analysis of TDLN.**
Detailed gating strategy used to analyse TDLN composition, including fluorescence-minus one control (FMO) to set the gate for AH1-specific cells.

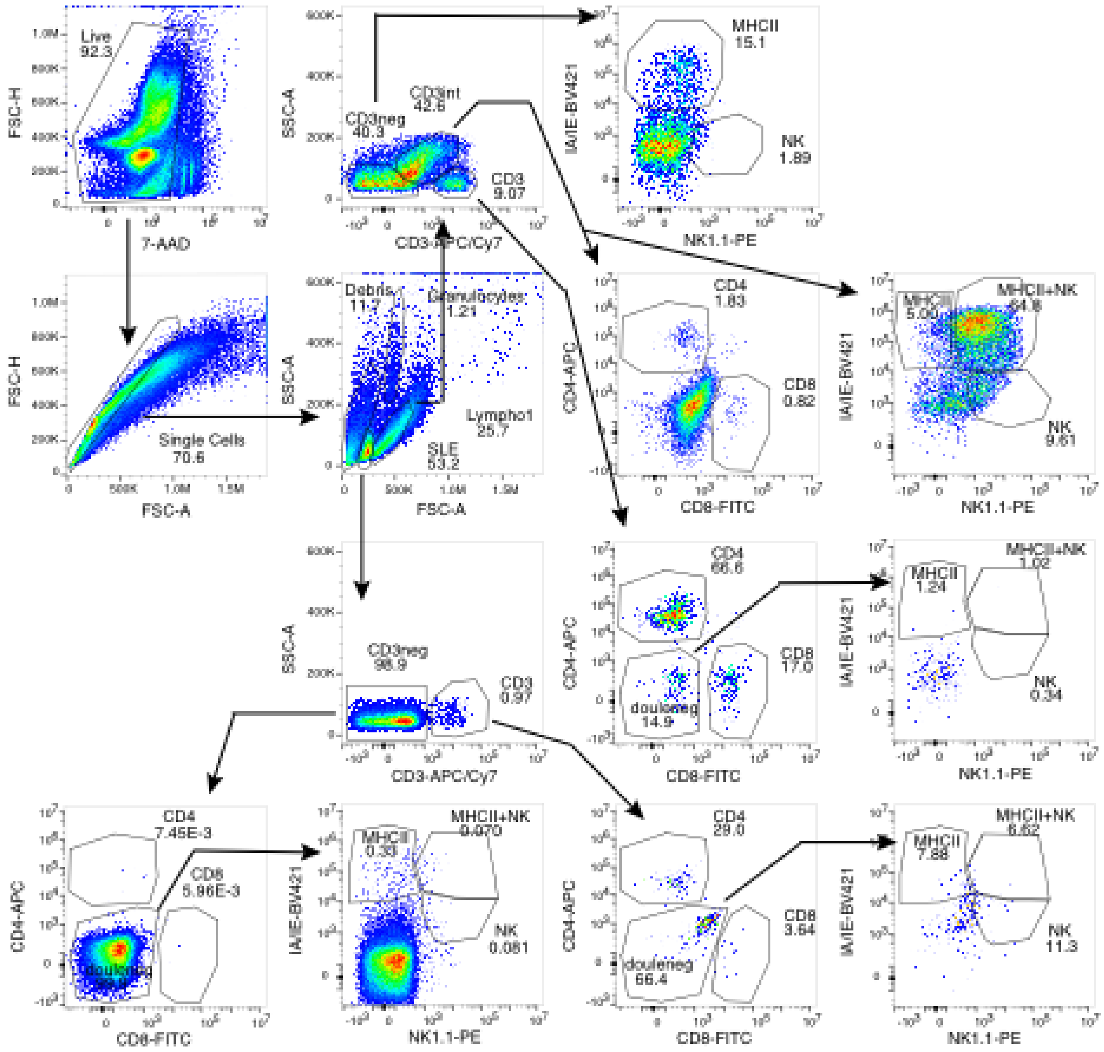


**Supplementary Figure 6. Gating strategy for analysis of tumor.**
Detailed gating strategy used to analyse tumor composition.

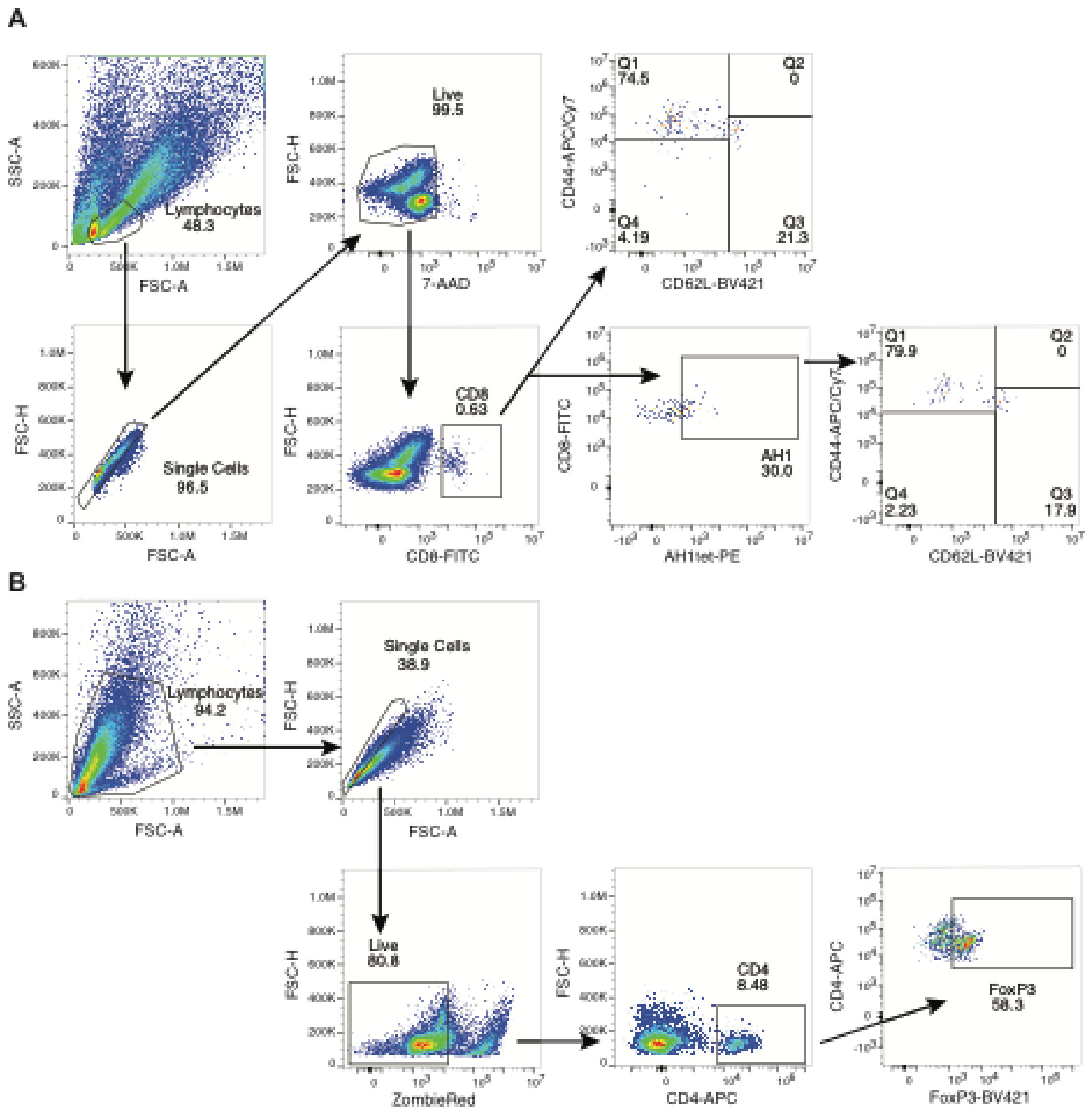


**Supplementary Figure 7. Gating strategy for tumor infiltrating CD8+ and CD4+ T cells.** Detailed gating strategy used to analyse AH1-specificity and phenotype of tumor-infiltrating $CD8^+$ T cells (**A**) and $CD4^+$ regulatory T cells ($T_{reg}$) (**B**). Gates for AH1, phenotype and FoxP3 were set using TDLN samples.
